# Alzheimer’s Disease Mutations Disrupt Neural Stem Cell Fate and Early Brain Development

**DOI:** 10.1101/2025.03.16.643546

**Authors:** Yiqiao Wang, Johan Lorentz, Elyas Mohammadi, Dilsah Ezgi Yilmaz, Theologos Sgouras, Yuxi Guo, Giusy Pizzirusso, Aphrodite Demetriou, Ivan Nalvarte, Marianne Schultzberg, Xiaofei Li

## Abstract

Alzheimer’s disease (AD) has been largely considered as an age-related disease, mainly affecting mature or aging adult brain. Recent studies show that AD-associated mutations could impair early life, even during neurodevelopment. However, due to the complex of AD mutations and neurodevelopmental regulations, how mutations in specific genes affect the origin of neurodevelopment is still largely under studied. In this study, we investigate how AD mutations in *App* gene impact neurodevelopment, with a focus on NSC dynamics and the balance between neurogenesis and gliogenesis. We employed the 5xFAD transgenic line and the APP^NL-G-F^ knock-in model, RNA sequencing, neurosphere assay and histological analyses on the cortex and hippocampus across critical developmental timepoints. Our results reveal that the APP^NL-G-F^ model exhibits early gene expression changes, with suppressed stem cell proliferation, impaired neurogenesis, upregulation of gliogenesis and enhanced neuroinflammatory pathways. In contrast, the 5xFAD model displays minimal embryonic differences, with pronounced postnatal alterations likely driven by both gene mutations and APP overexpression. These findings indicate that AD mutations can inherently impair NSC self-renewal and differentiation, resulting in a suboptimal brain structure that have potentially higher vulnerability towards AD pathology in later life.

## Introduction

The brain development is fundamentally orchestrated by neural stem cells (NSCs) residing in specialized niches of the embryonic brain, such as the subventricular zone (SVZ) and the hippocampus. These NSCs exhibit a delicate balance between self renewal and differentiation, which is crucial for generating the appropriate numbers and types of neurons, astrocytes, and oligodendrocytes needed to establish a properly structured and functional central nervous system^1^. Due to environmental insults or genetic mutations in critical regulatory genes, the disruption of this balance among different lineage differentiation can perturb the formation of neural circuits, rendering the developing brain more vulnerable to subsequent pathological insults and injuries. Such events have largely reported by many studies in the field of neural developmental disorders^2^. However, recent studies have begun to reveal that mutations associated with aging-associated disease, such as Alzheimer’s disease (AD) may also interfere with these fundamental developmental processes, predisposing the brain to neurodegeneration later in life^3–5^.

Alzheimer’s disease (AD) is classically understood as a neurodegenerative disorder mainly affecting the elderly, but growing evidence suggests its roots may extend to much earlier developmental stages^6–8^ (Ref). Supporting this notion, several genetic factors implicated in AD, including sporadic AD associated mutations ε4 allele of *APOE*, as well as familial mutations in *APP* and *PSEN1* genes could play critical roles in brain development and neural stem cell (NSC) biology. For example, the ε4 allele of *APOE*, the strongest genetic risk factor for sporadic AD, influences infant brain maturation: APOE4-carrying babies show altered myelination trajectories and reduced volumes in cortical regions vulnerable in AD^4,9^. These clinical and evolutionary perspectives highlight the importance of investigating how AD-associated genetic mutations impact the developing brain.

Familial AD (fAD) mutations in genes such as *APP* and *PSEN1* provide tractable models to examine neurodevelopmental consequences. *APP* is abundantly expressed in the embryonic and early postnatal brain and has essential functions in the balance of neuronal and glial differentiation, migration, and synapse formation^10^. Metabolites of APP (including soluble APPα) can regulate NSC proliferation and fate choice^11^, and disturbances in APP processing can therefore profoundly affect developmental neurogenesis^11^. Notably, excessive or misregulated APP signaling skews NSC fate: experiments in human neural stem cells have shown that elevating APP levels induces premature cell-cycle exit and pushes progenitors toward a glial lineage at the expense of neurons^12^ (REF). Conversely, knockdown of APP during corticogenesis leads to an accumulation of NPCs and disrupted migration, while APP overexpression in developing cortex drives cells to migrate past their appropriate cortical layer^13^. A proper balance of APP expression is therefore critical for normal neural proliferation, differentiation, and migration during development^10^. Familial AD mutations in *APP* (such as the Swedish, Arctic, and Iberian mutations used in mouse models) could perturb this balance by altering APP cleavage products (e.g. increasing amyloid-β or the APP intracellular domain), potentially disrupting the tightly regulated programs of neurogenesis and gliogenesis in the embryo and infant brain^14^.

*Presenilin-1 (PSEN1),* the catalytic subunit of γ-secretase, is likewise crucial for neurodevelopment. PSEN1 drives Notch receptor cleavage, a key signaling step in maintaining NSC populations. Complete loss of PSEN1 in mice causes embryonic lethal neurodevelopmental defects; conditional PSEN1 knockouts show premature differentiation of neural progenitors due to reduced Notch signaling (diminished Hes5 and elevated Dll1 levels)^15^. While heterozygous fAD mutations in *PSEN1* are compatible with development, they can subtly alter Notch signaling and other pathways. Indeed, recent studies using human cortical organoids with fAD mutations demonstrate that AD-linked *PSEN1* variants can aberrantly increase Notch activity during early development, leading to enlarged organoids with expanded progenitor pools and fewer differentiated neurons^6^. These findings reinforce that familial AD mutations are not silent until old age – rather, they can imprint the brain during developmental stages, potentially priming neural circuits for later vulnerability. Consistent with this, abnormal brain structural patterns have been observed in young adult carriers of fAD mutations decades before symptom onset^5^, and AD model organoids display distinct neurodevelopmental abnormalities depending on the specific mutation. In light of this emerging framework, we set out to investigate how AD-associated mutations impact early brain development, with a particular focus on neural stem and progenitor cell proliferation and differentiation. We employ two complementary AD mouse models: the well-established 5xFAD transgenic line (harboring five aggressive fAD mutations in *APP* and *PSEN1*)^16^ and the newer APP^NL-G-F^ knock-in line (carrying fAD mutations in the endogenous mouse App gene without overexpression)^17^. By examining these models at embryonic and early postnatal stages, we aim to disentangle the direct effects of AD mutations on neurodevelopmental processes from confounding factors such as transgene overexpression. We integrate RNA sequencing (RNAseq) analyses with cellular phenotyping and in vittro stem cell assays to assess how AD mutations alter NSC dynamics, neuronal differentiation, and glial development in the cortex and hippocampus during the peak periods of neurogenesis and gliogenesis. Situated within the broader context of AD’s neurodevelopmental implications, our study addresses a critical knowledge gap: How do the genetic drivers of AD mutations affect neurodevelopment? By framing AD mutations as not merely a cause for disease of aging but affect developmental origins, this work provides insight into early pathological mechanisms and lays groundwork for considering intervention long before the appearance of classical AD symptoms.

## Results

### AD mutations alter genetic regulations for stem cell property during development

To investigate whether AD-associated mutations affect embryonic and postnatal development, we perform bulk RNAseq on both cortex and hippocampus of 5xFAD, APP^NL-G-F^, and wildtype (WT) mice. Our primary focus is the selected timepoints for the peak of neurogenesis (E15.5), gliogenesis (E18.5) and postnatal development (P7). However, as no big differences were found between WT and 5xFAD during embryonic development (more details below), we also performed additional timepoints for these two mouse lines at P30 and P60. Comparing all datasets from the cortex and hippocampus samples in WT, 5xFAD and APP^NL-G-F^ mice at various developing timepoints, dimensional reduction analysis showed that APP^NL-G-F^ had more distinct features than the other two models from E15.5 to P7 (Fig 1B). To investigate the specific difference between animals carrying AD mutations compared to WT at each specific timepoint, we performed differential gene expression (DEG) analysis and revealed the most significant up- and down-regulated genes over time between genotype (Fig 1C-F). Consistently, in 5xFAD, very few DEGs were found during embryonic development (data not shown). From P7 on, we found that in 5xFAD mice, transcription factors associated with cell fate specification and neuronal maturation during brain development, such as *Scn4b* (P7 cortex), *Fgf3* (P30 cortex), and *Lhx9* (P60 cortex) were upregulated (Fig 1C), suggesting a potential pre-maturation of neurons due to the mutations in 5xFAD mice. We also observed subtle upregulations of immune response in the 5xFAD mice, for example *Cxcl10* was upregulated in the P60 5xFAD hippocampus (Fig 1D). Interestingly, while most of the DEGs did not show high fold changes between 5xFAD and WT mice (Fig 1C-D), the most significantly upregulated genes in 5xFAD brains were *Psen1*, *Thy1* and *App* (P60 in Fig 1C-D), suggesting that the alterations could be due to the synergic effects of these AD-associated transgenes, but might be difficult to pinpoint the effects of individual gene of these three.

**Figure 1.**
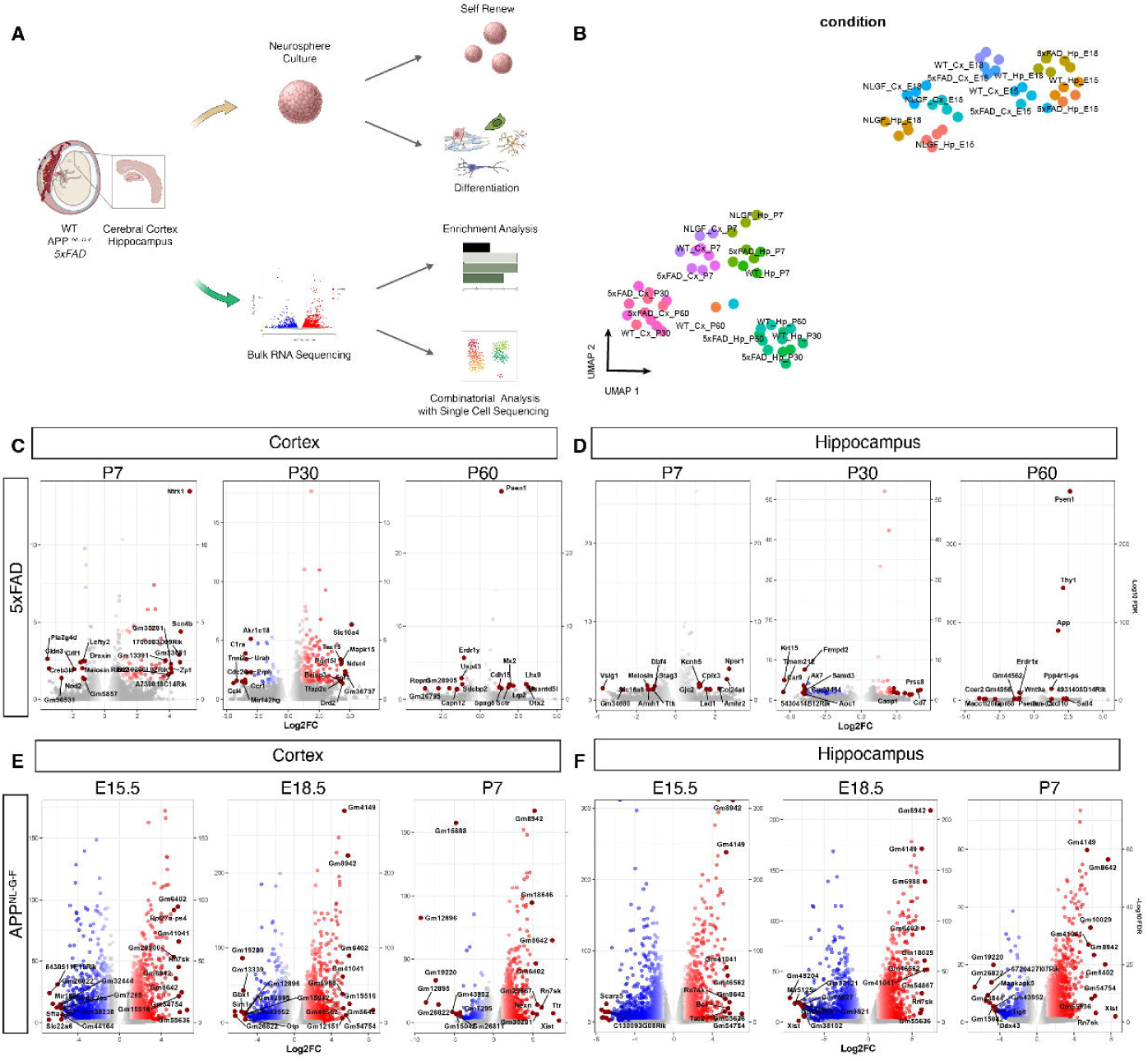
AD mutations alter gene expression programs underlying stem cell properties during development. **(A)** Schematic illustration of the experimental design. Embryonic (E15.5, E18.5) and postnatal (P7, P30, P60) cortices and hippocampi from wild-type (WT), 5xFAD, and APP^NL-G-F^ mice were collected for neurosphere assays, immunostaining, bulk RNA-sequencing (RNA-seq), and association with scRNAseq. **(B)** Dimensional reduction analysis (UMAP) of all RNA-seq samples, showing that APP^NL-G-F^ clusters distinctly from 5xFAD and WT from E15.5 to P7. Each dot represents an individual sample; color denotes genotype. **(C, D)** Volcano plots of differentially expressed genes (DEGs) comparing 5xFAD vs. WT at indicated timepoints in cortex (**C**) and hippocampus (**D**). Top DEGs are labeled. Upregulated genes are shown in red, downregulated genes in blue; non-significant genes are in gray. **(E, F)** Volcano plots of DEGs comparing APP^NL-G-F^ vs. WT at indicated timepoints in cortex (**E**) and hippocampus (**F**).

In contrast, in the APP^NL-G-F^ mice, we observed more robust DEGs at much earlier stages, from E15.5-P7 (Fig 1E-F). We found upregulation of genes associated with NSC fate specification, specifically to gliogenesis such as *Sox10* and *Gfap*, as well as potential early neuroinflammation with higher expression in *Cxcl10*. In contrast, genes related to neuronal subtype specification were downregulated, such as *Gbx1* and *Sim1* for motor neuron or serotonergic specification^18^. Notably, due to large number of DEGs were found in the analyzed outcome, not all of them were highlighted in the volcano plots in Fig 1. The full list of DEGs in each condition can be seen in table 1. Our data suggest that AD mutations could disturb stem cell property during brain development.

### 5xFAD perturb neurodevelopment via AD mutations and App overexpression

To further investigate how different AD pathology mouse models affect brain development, we first performed gene enrichment analysis on the DEGs in 5xFAD mice compared to WT from P7 to P60 (Fig 2A-B). We found that in 5xFAD cortex, synaptic transmission of GABAergenic and cholinergic neurons and axon guidance were upregulated throughout these three timepoints (Fig 2A). In the 5xFAD hippocampus, besides axongenesis and neurotransmitter secretion, we also found that glial cell activation, astrocyte differentiation and neuroinflammatory response were upregulated (Fig 2B). These data suggest that there could be an overall faster pace of stem cell differentiation and neuronal migration at early life of the 5xFAD mice, at the expense of stem cell niche and a proper neural network establishment. However, such events could have been shown earlier during embryonic development. To understand why the difference at embryonic developmental stages was subtle, and whether such disturbed postnatal development was solely caused by the AD mutation on *App* and *Psen1* gene, we plot the gene expression of *Thy1*, *App* and *Psen1* from all our datasets in 5xFAD mice, since the *App* and *Psen1* genes were expressed under the *Thy1* promoter. We found that *Thy1* was not expressed until postnatal in either the cortex or the hippocampus (Fig 2C), suggesting that the mutations in *App* and *Psen1* were not present during embryonic stages either. The low expression of *App* before birth is therefore the endogenous *App* gene (Fig 2C), which started increasing postnatally, in line with the expression of *Thy1*. This explains the lack of difference between 5xFAD and WT from E15.5 to E18.5. In postnatal timepoints, we observed higher expression of these three genes in 5xFAD samples from both cortex and hippocampus compared to WT (Fig 1C), in line with previous studies showing App protein is over produced in 5xFAD mice^17^. Moreover, since *App* is also important for stem cell self-renewal and differentiation^10,11,19^, we further studied whether the postnatal differences was due to the combination of gene mutations and App expression level. Indeed, using the 5xFAD cortex as an example, our gene correlation analysis confirmed that the top upregulated DEGs in 5xFAD mice were all positively correlated with the expression level of *Thy1* and *App*. These data suggest that the developmental disturbance in postnatal 5xFAD mice is largely contributed by the synergy of AD mutations and the over expression of *App* and *Psen1*. Due to the complex contributors to the phenotype we had observed in 5xFAD mice, we did not continue further investigation in this mouse model for brain development.

**Figure 2.**
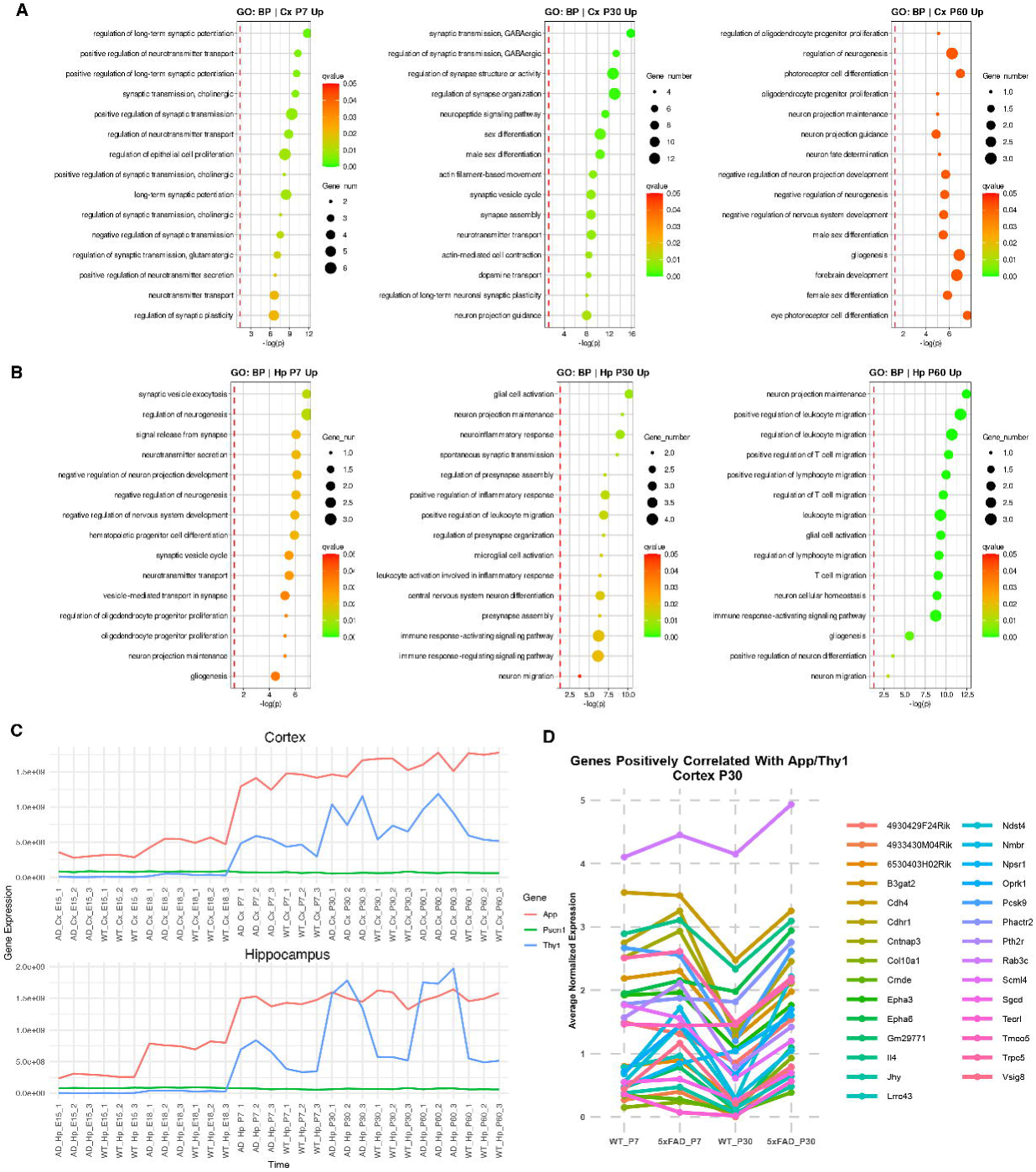
5xFAD perturbs neurodevelopment via synergy of AD mutations and *App* overexpression. **(A, B)** Gene Ontology (GO) enrichment analysis of DEGs in 5xFAD cortex (**A**) and hippocampus (**B**) relative to WT from P7 to P60. Dot size indicates the number of genes associated with each GO term; color scale indicates the adjusted *p*-value (dark orange = more significant). Enriched pathways include synaptic transmission, axon guidance, glial cell activation, and inflammatory response, suggesting accelerated or dysregulated developmental processes in 5xFAD. **(C)** Expression profiles of *Thy1*, *App*, and *Psen1* across embryonic and postnatal timepoints in 5xFAD cortex and hippocampus. **(D)** Correlation analysis between Thy1/*App* expression and top upregulated DEGs in the 5xFAD cortex (P30). DEGs with high correlation to *Thy1* and *App* are selected, indicating that overexpression of mutant *App* and *Psen1* is a major driver of postnatal transcriptional alterations.

### AD mutations in App gene alone contribute to imbalanced neurogenesis and gliogenesis

To pinpoint how AD mutations alone could affect early neurodevelopment, we used the APP^NL-G-F^ mice, in which the AD mutations were knocked in the endogenous App gene without causing App overexpression^17^. We first showed that the *App* expression started as early as E15.5, and confirmed no significant difference between APP^NL-G-F^ and WT mice (Fig 3A). We performed gene enrichment analysis on the DEGs from APP^NL-G-F^ mice in the cortex, and found that synaptic formation, neuronal apoptotic process, glial cell activation and activation of microglia and immune cells were upregulated across all these time points (Fig 3B). These data suggest that there could be a pre-maturation of the developing brain of APP^NL-G-F^ mice. To analyze how these gene expression alteration affects neurodevelopment at the cell level, we focused on NSCs and their progeny, as well as microglial development by immunostaining. We found that there are no significant differences of the total number of NSCs (Sox9+) in the stem cell niche (subventricular zone, SVZ) between APP^NL-G-F^ and WT mice from E13.5 to P7 (Fig 3C-D). However, the proliferation capacity of NSCs (Sox9+Ki67+) in APP^NL-G-F^ mice was significant lower at E15.5 and E18.5, at 34.2% and 20.8%, respectively, compared to 44.2% and 20.8% in WT. By evaluating the neuronal differentiation at the end point of our investigation at P7, we found that there was a significantly lower number of mature neurons in the APP^NL-G-F^ cortex (60.83%) compared to WT (73.67%) at P7 (Fig 3E-F). Our data suggest that AD mutations in the *App* gene suppress stem cell proliferation and neuronal differentiation during development.

**Figure 3.**
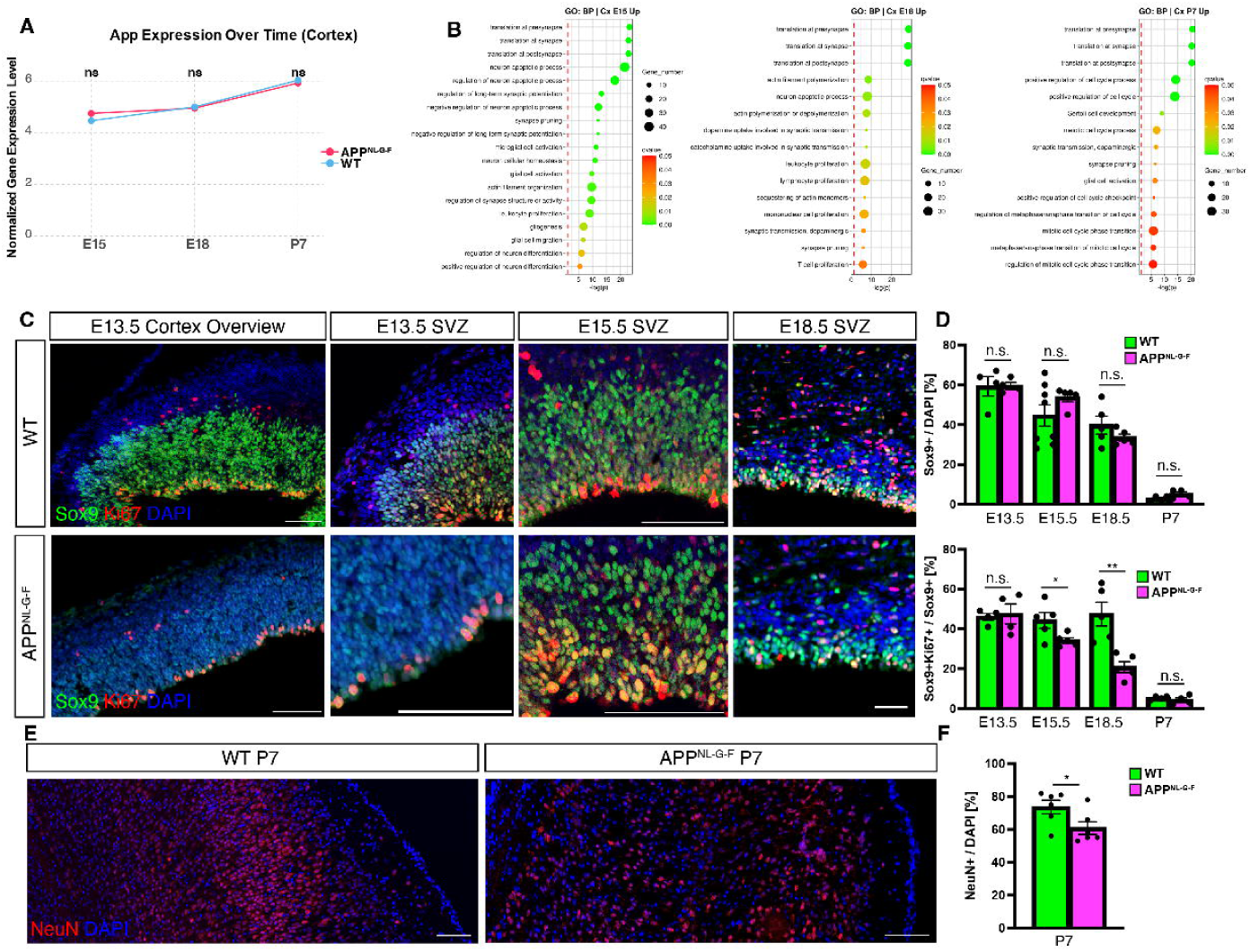
AD mutations in *App* alone contribute to imbalanced neurogenesis and gliogenesis in the cortex. **(A)** *App* expression in WT and APP^NL-G-F^ cortices from E15.5 to P7, measured by RNA-seq. No significant difference in overall *App* levels is observed between genotypes. **(B)** GO enrichment analysis of DEGs in APP^NL-G-F^ cortex vs. WT, highlighting upregulated pathways associated with synapse formation, neuronal apoptosis, glial activation, and immune responses across E15.5, E18.5, and P7. **(C)** Representative immunofluorescence images of Sox9+ cells (green), Ki67+ cells (red) and total cells (DAPI, blue) atindicating stages. **(D)** Quantification of indicating marker positive cells and their ratio. **(E)** Representative images of mature neurons (NeuN+, red) at P7 in WT and APP^NL-G-F^ cortices. **(F)** Quantification of NeuN+ cells reveals a significant reduction in neuronal differentiation in APP^NL-G-F^ mice compared to WT at P7. Error bars represent mean ± s.e.m. (*p* < 0.05).

Since the GO terms in RNAseq also revealed upregulation of gliogenesis and mímmune cell activation in the APP^NL-G-F^ mice (Fig 1C and Fig 2A), to study how such mutations affect gliogenesis, we performed immunostaining of OPC marker Olig2 and developing astrocyte marker Aldh1l1 at various timepoints in the cortex. We observed limited number of OPCs from E13.5-15 during their first wave generation for sufficient comparison, thus we turned to the lateral ganglionic eminence (LGE), the niche of OPCs at this stage. While no significant difference was found at E13.5 (Fig 4A-D, I), there was a significantly higher number of Ki67+ OPCs between genotypes at E15.5 (Fig 4E-I), suggesting a larger oligodendrocyte differentiation potential of OPCs in the APP^NL-G-F^ mice. With the development progresses, OPCs migrate to the cortex and continue development. Therefore, we quantified the ratio of proliferative OPCs in the cortex and found significantly higher number of Ki67+ OPCs at both E18.5 and P7 in the APP^NL-G-F^ cortex (41.50% and 3.25%, respectively), compared to those in the WT cortex (23.33% and 0.67%, respectively) (Fig 4J-R). While the signals of astrocyte marker Aldh1l1 at embryonic stages were not sufficient to compare these two mouse lines, we observed a significant difference at P7, with higher number of astrocytes in the cortex in APP^NL-G-F^ mice with 11.47%, compared to WT with 8.98% (Fig 4S,T,W). These data suggest that AD mutations in *App* gene promote gliogenesis during development, probably at the expense of neurogenesis (Fig 3E-F). Interestingly, as our RNAseq data revealed upregulated DEGs related to microglial activation at E18.5 and P7 (Fig 3B), we performed Iba1 staining on the tissue sections to assess the microglia at early development. While there was no difference between genotypes at E15.5 and E18.5 (data not shown), there was a more than one-fold increase of Iba1+ cells in the APP^NL-G-F^ cortex compared to WT (6.80% *v.s.* 3.00%). As higher expression of immune-related genes was shown in our RNAseq data (Fig 1E, Table 1), our results here suggest that AD mutations on *App* gene could lead to earlier microglia activation and neuroinflammation at early life.

**Figure 4.**
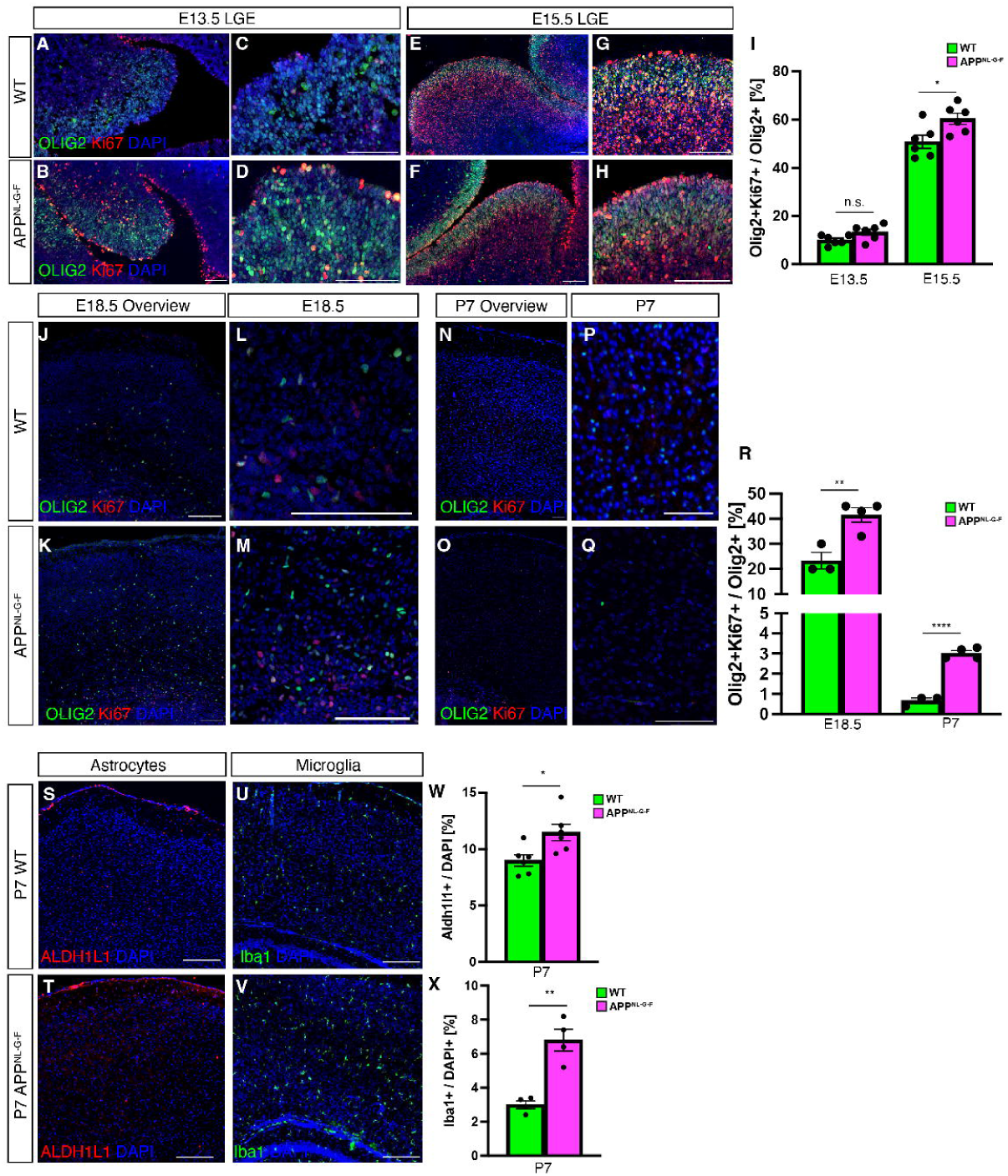
AD mutations in *App* enhance gliogenesis and induce early microglial activation in the cortex. **(A–H)** Representative immunostaining of Olig2 (green) and Ki67 (red) in the lateral ganglionic eminence (LGE) at E13.5 (**A–D**) and E15.5 (**E–H**) for WT and APP^NL-G-F^ mice. C, D, G, H are in higher magnificantion to A, B, E, F, respetively. Scale bars, 100 µm. **(I)** Quantification of proliferating (Ki67+) OPCs. **(J–O)** Olig2 (green)/Ki67 (red) staining in the cortex at E18.5 (**J–M**) and P7 (**N–O**). L, M, P, O are in higher magnification to J, K, N, O, respetively. **(R)** Quantification of proliferating OPCs (Ki67+ Olig2+) indicates increased OPC proliferation in APP^NL-G-F^ compared to WT at both E18.5 and P7. **(S-V)** Astrocyte marker Aldh1l1 (red) immunostaining (S, T), and microglia marker staining (U, V) at P7 in WT and APP^NL-G-F^ cortices. **(W-X)** Quantification of Aldh1l1+ astrocytes (W) and microglia (X).

To investigate whether AD mutations also affect the other stem cell niche, the hippocampus, we performed similar analysis of GO and immunostaining (Fig 5). Similarly, we first confirmed that *App* expression is at the same level between genotypes (Fig 5A). With GO analysis, we found that the upregulated DEGs in APP^NL-G-F^ mice were related to cell cycle process, synaptic formation, neuronal apoptosis, gliogenesis and immune response (Fig 5B), similar to thoese terms in the cortex (Fig 3B). Our immunostaining results showed that total Sox9+ cells in the developing hippocampus from E13.5 to E18.5 had no significant differences between genotypes (Fig 5C, E), but the proliferative potential was significantly lower in APP^NL-G-F^ hippocampus, shown by the ratio of Ki67+Sox9+ cells (Fig 5C, F). Notably, the differences at E13.5 was dramatic (Fig 5F), which might need additional animals to confirm in future work. Besides the decreased proliferative potential in in the APP^NL-G-F^ hippocampus, we also found early microglial activation in the APP^NL-G-F^ hippocampus at P7, as there was substantial higher Iba1+ cells compared to WT (Fig 5D, G), similar to the findings in the cortex. Therefore, our data suggest that AD mutations in the *App* gene impair braian development via suppressing NSC proliferation and neuronal differentiation, promoting excessive glial differentiation, and inducing early microglial activation and potential neuroinflammation.

**Figure 5.**
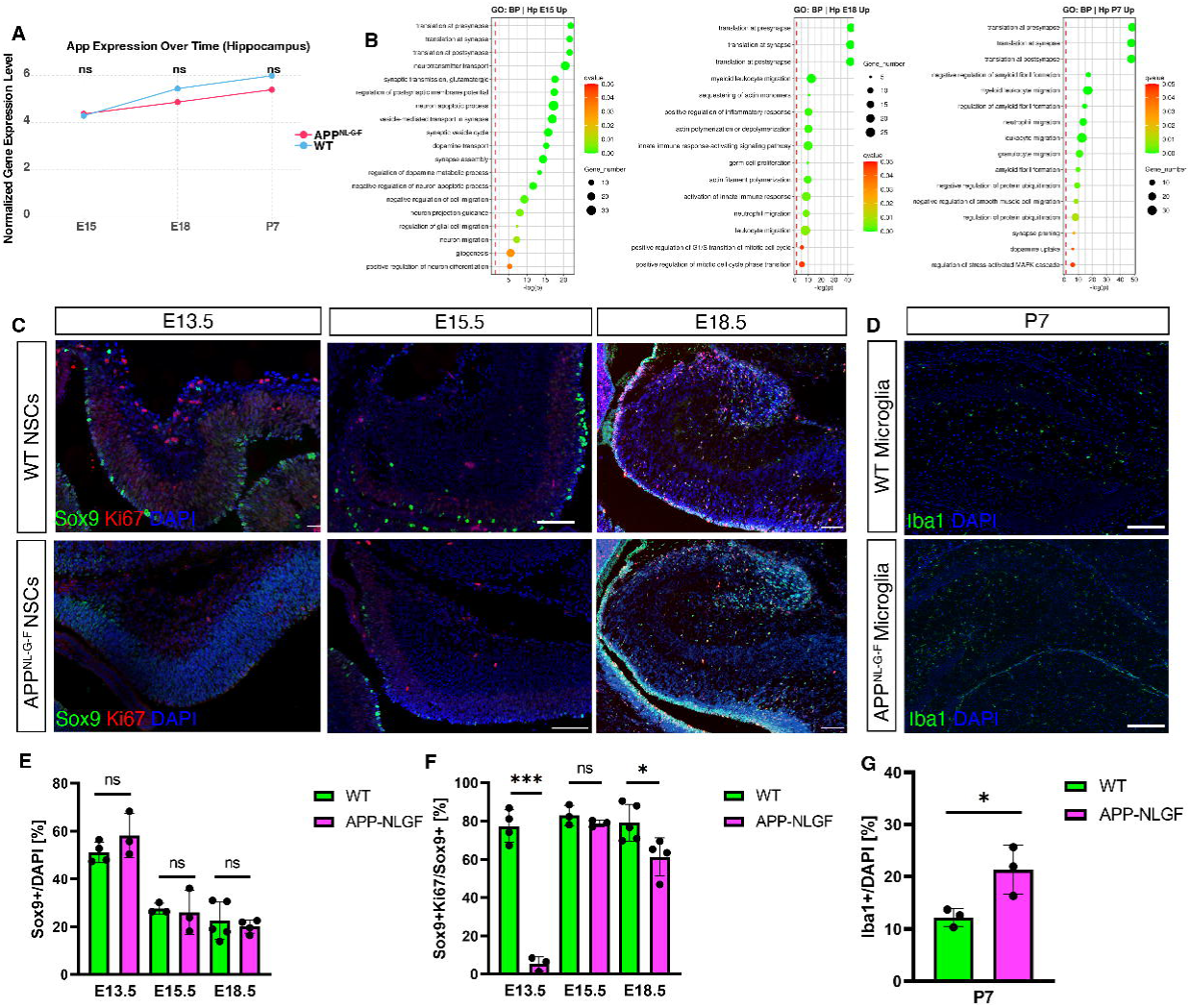
AD mutations in *App* disrupt hippocampal development. **(A)** *App* expression in WT and APP^NL-G-F^ hippocampi from E13.5 to P7, confirming no overexpression in the knock-in line. **(B)** GO enrichment analysis of DEGs in APP^NL-G-F^ hippocampus vs. WT highlights upregulated pathways for cell cycle, synaptic formation, neuronal apoptosis, gliogenesis, and immune response across E13.5 to P7. **(C)** Representative images of Sox9 (green)/Ki67 (red) staining in WT and APP^NL-G-F^ hippocampi at E13.5, E15.5 and E18.5. **(D)** Representative Iba1+ microglia staining at P7 in WT vs. APP^NL-G-F^ hippocampus. **(E-G)** Quantification of total Sox9+ NSCs, Ki67+Sox9+ NSCs and microglia in the cortex of indicating timepoints.

### AD mutations in App disrupts self-renewal and the balance of neuronal and glial differentiation in vitro

To further investigate how the AD mutations affect the intrinsic stem cell potential of NSCs, we performed neurosphere assay to assess the self-renewal potential in WT and APP^NL-G-F^ mice during development. We dissected the embryonic cortical regions from E13.5 and 18 from both genotypes, and grew neurospheres for 1 week *in vitro*. Consistently across all timepoints, we found that the number of neurospheres was significantly lower from the APP^NL-G-F^ mice compared to those from WT, and this difference continued after passage at both E13.5 (Fig 6A-B) and E18.5 (Fig 6C-D). After passage, neurospheres are supposed to derive from only NSCs as neural progenitor-derived neurospheres has limited passage ability^20^. Our data showed that the there was a substantial loss of neurospheres generated in the second generation of neurospheres at both E13.5 and E18.5, suggesting that *App* mutations impair the self-renewal potential of NSCs. To further evaluate the differentiation capacity of these cultured stem cells, we differentiated the primary neurospheres 14 days *in vitro* from the E13.5 primary neurospheres (Fig 6B and D). In APP^NL-G-F^ neurospheres, the capacity to generate neurons is significantly decreased as shown by less Tuj1+ cells, in line with the in vivo data. Similarly, the ratio of differentiated GFAP+ cells was significantly higher in APP^NL-G-F^, supporting the in vivo analysis of higher astrocytes generated in the APP^NL-G-F^ cortex (Fig 6E-F). Interestingly, the ratio of CNPase+ OPCs was significantly lower from the APP^NL-G-F^ group, opposite to the observation from *in vivo* (Fig 4). This is probably due to AD mutations in *App* gene affecting different source of OPCs differently, which may suppress OPC differentiation from NSCs while promote OPC differentiation from fate-committed precursors of OPCs. This requires further investigation by cell-type specific analysis. Taken together, our in vitro data suggest that AD mutations in *App* gene suppress NSC self-renewal, promote astrocyte differentiation, and inhibit neuronal and OPC differentiation.

**Figure 6.**
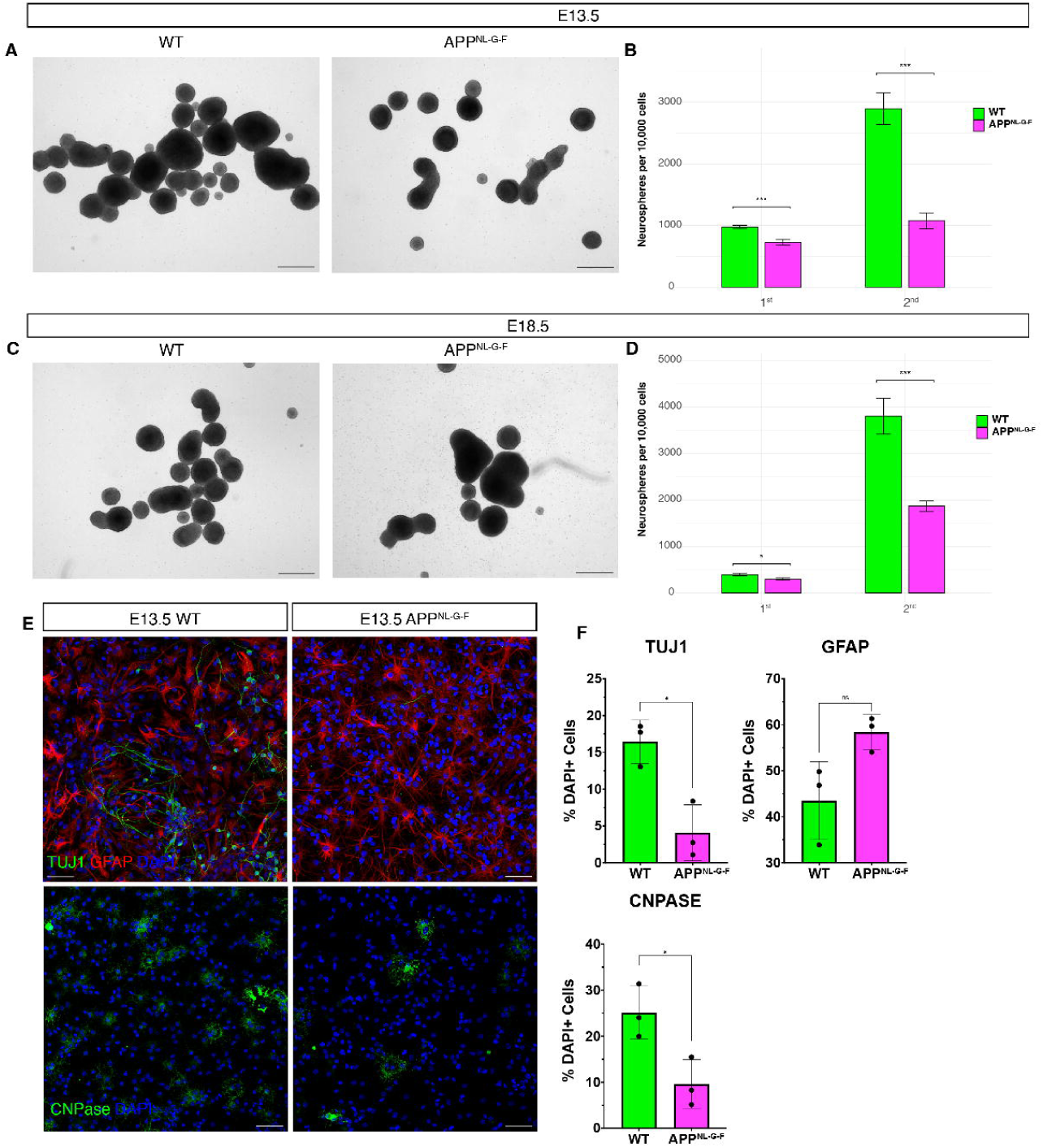
AD mutations in *App* impair NSC self-renewal and alter differentiation fates in vitro. **(A, C)** Representative images of neurospheres derived from E13.5 (**A**) and E18.5 (**C**) WT and APP^NL-G-F^ forebrain cultured for 7 days in vitro. Scale bars, 200 µm. **(B, D)** Quantification of neurosphere number per 100,000 cells in primary (1^st^) and secondary (2^nd^) cultures. **(E)** Immunofluorescence staining of differentiated neurospheres from E13.5 cortical cultures (14 days *in vitro*) for TUJ1 (neurons, red), GFAP (astrocytes, green), and CNPase (oligodendrocytes, green). **(F)** Quantification of lineage markers. Error bars represent mean ± s.e.m.

### Characterization of heterogenous cell types affected by AD mutations during development

Since the developing brain has heterogenous cell types, we wonder whether the *App* mutations mainly affect specific cell type. Therefore, we leveraged the information from a single-cell RNA sequencing (scRNAseq) developmental cell atlas of mouse cortex (Fig 7A-B) of the same genetic background (c57 mice)^21^. We investigated the shared DEGs between scRNAseq and bulk data to identify where/when the gene deregulation is taking place. Hence, this intersection analysis was done for cell types-specific and time-point-specific DEGs in scRNAseq dataset separately (Fig 7 C-F). We found that among the top up and downregulated DEGs, there was a cell type specific gene expression pattern. For example, upregulated DEGs in the APP^NL-G-F^ mice, such as *Scrg1*, which has shown to be associated with OPC differentiation and myelination^22^, has specific expression in oligodendrocytes (Fig 7C). This partly explained the earlier oligodendrocyte differentiation in our study. Moreover, *Ccl12* and *Lst1* with direct roles in microglia development and proinflammation functions^23,24^, showed exclusive expression in microglia (Fig 7C). This further provided more insights in the mechanisms of our observation of higher number of microglia in both APP^NL-G-F^ cortex and hippocampus, alongside with the upregulated genes associated with inflammation. In addition, downregulated gene *Cldn11* in our RNAseq data was shown specifically expressed in vascular leptomeningeal cells (VLMC) during cortical development (Fig 7D). As *Cldn11* contribute to the specialized functions of endothelial transport and tight junction formation, such alteration suggests that AD mutations in the *App* gene affect not only the neural lineage but also the permeability and homeostatic functions of the blood brain barrier, potentially leading to developmental and functional disturbances in the brain. Finally, we also surveyed whether the DEGs revealed in our RNAseq data also have a timepoint specific expression pattern. Among the top DEGs with cell-type specific expression pattern, we found some of them also exhibit distinct expression at different developmental stages (Fig 7E-F). Collectively, our results show that AD mutations have broad effects on various cell types during brain development, which could lead to an abnormal structure that is more vulnerable to future pathology, such as Aβ plaques and tau tangles in later life.

**Figure 7.**
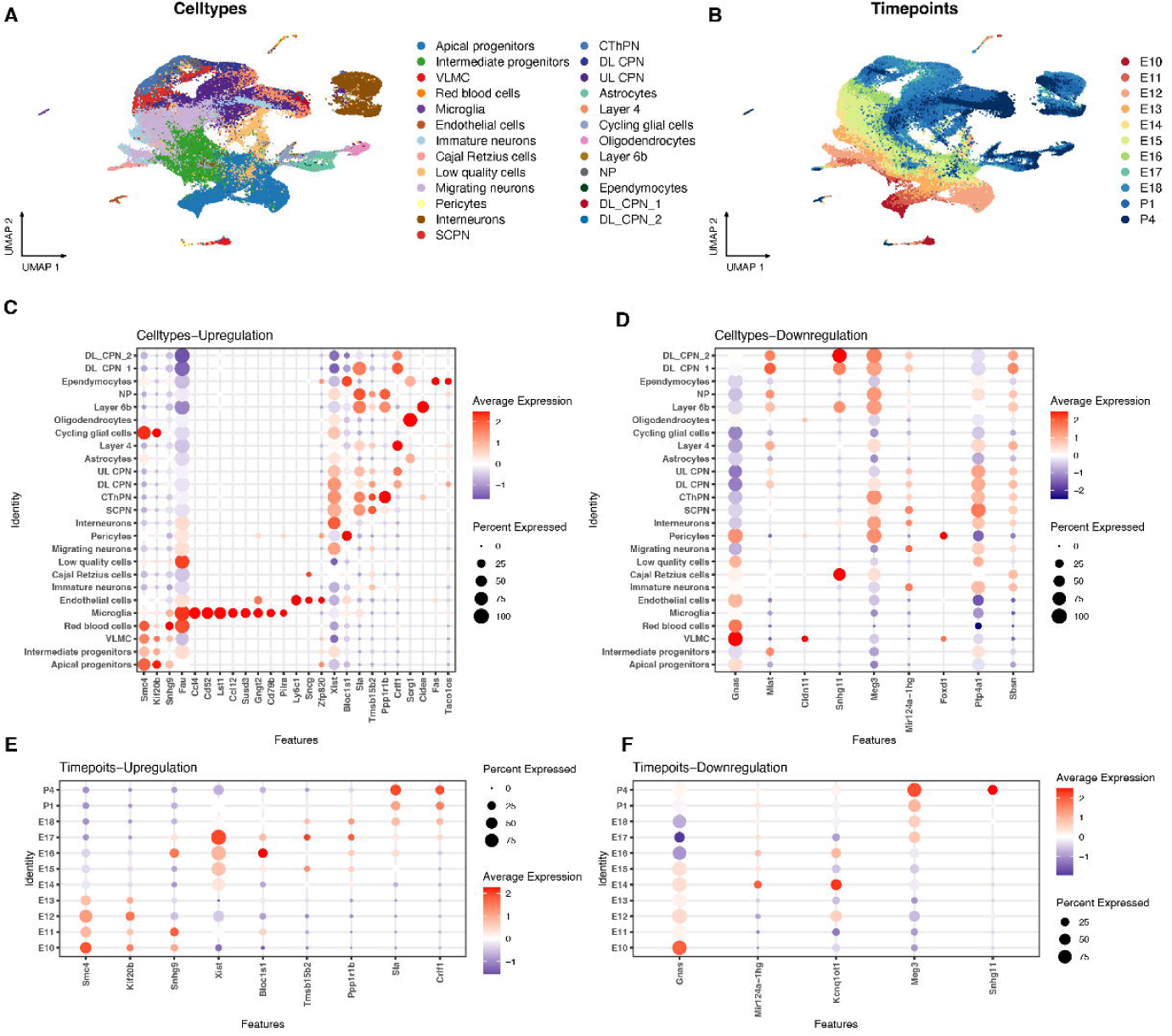
Single-cell atlas analysis reveals cell-type- and timepoint-specific impacts of AD mutations. **(A, B)** UMAP plots from a developmental scRNA-seq atlas of WT mouse cortex, showing major cell populations (**A**) and timepoints (**B**) across embryonic and early postnatal stages. **(C, D)** Dot plots indicating average expression (color) and percentage of cells expressing each gene across relevant cell types for upregulated (**C**) and downregulated (**D**) DEGs in APP^NL-G-F^. **(E, F)** Dot plots showing expression of key DEGs across different developmental timepoints in the scRNA-seq atlas, by using up- and down-regeulated DEGs in APP^NL-G-F^ mice.

## Discussion

Our findings demonstrate that i) familial AD mutations can disrupt early brain development by affecting NSC proliferation and the balance between neurogenesis and gliogenesis; ii) the expression of key transcription factors related to cell proliferation, neuronal and glial differentiation, as well as and psynatic formations were perturbed; and iii) microglia show early activation during development and early inflammation under the influence of familial AD mutations. Notebly, in interpreting whether or not certain AD mutations affect early brain development, it is crucial to consider the animal models used, such as whether the key genes carrying mutations have altered expression level. Below, we discuss the implications of our results with respect to model suitability, consistency with prior studies, translational relevance, and remaining limitations.

### 5xFAD model: AD mutations vs. APP overexpression in development

The 5xFAD mouse is a widely used model that recapitulates key aspects of AD amyloid pathology at relatively young ages (developing robust Aβ deposition and neuroinflammation by a few months of age)^25^. However, 5xFAD carries human APP and PSEN1 transgenes under the Thy1 promoter, which substantially complicates its use in developmental studies. The Thy1 promoter is largely inactive during embryogenesis and only becomes active around one week after birth^26^. Consistent with this, we observed no significant transcriptional or cellular differences in 5xFAD embryos compared to wild-type, and only postnatal time points (P7 and onward) began to show divergence. This timing aligns with the delayed expression of the mutant transgenes: our data showed that *Thy1* itself (the driver of the transgenes) is not expressed in cortex or hippocampus until the postnatal period, explaining the lack of embryonic phenotype in 5xFAD. Once Thy1-driven expression initiates, 5xFAD brains begin to overexpress mutant *APP* and *PSEN1*, leading to an acceleration of developmental processes such as neuronal differentiation and migration in early life. We found upregulation of pathways related to synaptogenesis, axon guidance, and early gliogenesis in young 5xFAD brains, suggesting a precocious maturation of neural elements. Notably, many of the gene expression changes in postnatal 5xFAD mice were correlated with *APP* levels, implying that the high dosage of APP (and its increased cleavage products) is a major contributor to the phenotype. This highlighhts a key caveat: 5xFAD’s neurodevelopmental effects arise from a combination of AD mutations and overexpression, rather than mutations alone. Overexpression of APP is itself known to perturb neurodevelopment – for instance, raising APP levels in neural progenitors drives them toward glial fates and reduces neuronal output^12^.

Our results in 5xFAD are congruent with this: by P30-P60, 5xFAD mice showed evidence of an exhausted stem cell niche and heightened glial/immune activation, as would be expected if excessive APP signaling had accelerated stem cell differentiation at the expense of self-renewal. Thus, while 5xFAD is useful for modeling aggressive amyloid pathology, its developmental confounds (non-physiological expression pattern and gene dosage) mean it must be interpreted with caution in the context of early brain development^17^. The 5xFAD model may exaggerate or prematurely induce developmental phenotypes that are not purely mutation-driven, highlighting the importance of complementary models that more faithfully mimic endogenous gene expression.

### AD mutations in App alone are sufficient to alter neurodevelopmental trajectories

To isolate the effect of AD mutations without overexpression artifacts, we turned to the APP^NL-G-F^ knock-in model, and our data show that AD mutations in App alone are sufficient to alter neurodevelopmental trajectories. In APP^NL-G-F^ embryos and neonates, where mutant App is expressed at physiological levels from early stages, we detected clear deficits in NSC proliferation and neuronal differentiation that were absent in 5xFAD at the same ages. These knock-in mice had fewer proliferating Sox9+ neural stem/progenitor cells at E15.5–E18.5 and generated fewer neurons by P7, despite having a normal total pool of NSCs. Concomitantly, they exhibited increased oligodendrocyte progenitor (Olig2+) proliferation and elevated astroglial (Aldh1l1+) and microglial (Iba1+) populations in the early postnatal cortex. In essence, the App mutations biased developing cells away from neurogenesis and toward gliogenesis, mirroring in vivo what had been observed *in vitro* when APP levels or amyloid-β signaling are high^12^. These findings resonate with other AD models. Notably, the 3xTg-AD mouse (harboring mutant APP_Swe, PSEN1_M146V, and tau_P301L) also displays early-life neurogenic impairments: 3xTg-AD mice have significantly reduced hippocampal stem/progenitor proliferation and newborn neurons as early as postnatal day 7, well before amyloid plaques or tau tangles form^8^. That study also reported smaller hippocampal volumes and altered expression of developmental regulators (e.g. Notch and Wnt pathway genes) in one-month-old 3xTg mice^8^, consistent with a developmentally derailed neurogenic program. Our APP^NL-G-F^ model, which isolates mutant APP’s contribution, produces a similar theme of early NSC dysfunction, strengthening the evidence that familial AD-linked genes intrinsically perturb brain development across different model systems. It is worth noting differences among models as well: for example, organoid models of *PSEN1* mutations have found an increase in progenitor proliferation with delayed neuronal differentiation (attributed to heightened Notch signaling)^6^, whereas our APP mutant mice showed reduced progenitor proliferation. These discrepancies likely reflect distinct mechanistic effects of APP vs. PSEN1 mutations – the former may accelerate progenitor cell-cycle exit via toxic APP fragments or altered β/γ-cleavage products, whereas the latter may prolong progenitor maintenance via Notch pathway alterations^6^. In both cases, however, the end result is an imbalance in the normal progression of neurogenesis, reinforcing the concept that multiple AD-related mutations, through different pathways, can disrupt the timing and quality of early brain development.

Beyond 5xFAD and APP^NL-G-F^, other AD mouse models provide additional context. Traditional APP overexpression models (e.g. PDGF-hAPP or Thy1-APP transgenics) have long been noted to cause developmental abnormalities in the absence of any mutant allele; for instance, Naumann et al. reported that overexpressing wild-type human APP in a mouse hippocampus impaired the activity-dependent maturation of newborn neurons^27^. Similarly, APOE4 knock-in mice (while not a model of familial AD, they model a risk factor) exhibit synaptic and cognitive alterations early in life, presumably due to developmental effects of the E4 allele on neurons and glia (Ref). On the other hand, mice modeling solely *PSEN1* mutations (without *APP* changes) have been less extensively studied in development because many *PSEN1* mutations do not cause overt phenotypes in young mice. However, subtle molecular changes have been observed. For example, knock-in mice with certain PSEN1 fAD variants show altered adult neurogenesis and synaptic activity later on^28,29^, hinting that developmental perturbations might have occurred. In humans, it remains challenging to directly observe neurodevelopmental consequences of fAD mutations, since carriers typically appear cognitively normal until mid-adulthood. Nonetheless, structural MRI studies of at-risk individuals lend support to early changes – young asymptomatic carriers of AD mutations or APOE4 often show differences in gray matter volume or network connectivity compared to controls^30,31^. Thus, while animal and stem cell models each have idiosyncrasies, a convergent picture emerges: AD-related genetic alterations can modulate fundamental developmental processes like stem cell proliferation, differentiation, and brain circuit formation.

### Translational implications in human patients

If AD mutations indeed skew neural development, what might this mean for human disease? One possibility is that developmental dysregulation creates a brain that is subtly anatomically and functionally altered from the start – for example, an imbalance in excitatory/inhibitory neuron production, or a reduced reserve of neural stem cells, or a primed pro-inflammatory state due to early microglial activation. These changes might not cause obvious deficits in childhood, due to the remarkable plasticity and redundancy of the young brain, but they could render certain neural networks more vulnerable to aging or to a second “hit” later in life. In our APP^NL-G-F^ mice, the increased glial cells and activated microglia observed at P7 suggest that AD mutations create an inflammatory milieu even in early life. Translationally, this is intriguing given that neuroinflammatory markers (e.g. activated microglia via PET imaging or cytokines in CSF) have been detected in people decades before they develop AD symptoms, including in presymptomatic familial AD cases. Early excessive gliogenesis at the expense of neurogenesis, as we saw in mutant APP mice, could potentially contribute to a form of developmental gliosis or altered brain connectivity. In a human context, one could speculate that such early alterations might manifest as subtle differences in cognitive development or behavior, although this remains unproven. More concretely, recognizing a developmental component to AD opens the door to consider very early interventions. For example, if heightened Notch signaling or Wnt dysregulation is identified as a consequence of an AD mutation during gestation or infancy, one could envision targeted therapies or environmental interventions applied in early life to normalize developmental trajectories, long before amyloid plaques appear^6^. While treating embryos or infants for a late-life disease is far from current clinical practice, the concept raises a shift in thinking: prevention of AD may need to start as early as neurodevelopment, at least from juvenile stage for the clinical feasibility, especially for those with known high-risk mutations. Additionally, our study highlights the relevance of using human-model systems (such as induced pluripotent stem cells) alongside animal models to confirm which developmental phenotypes translate across species. The cortical sphere models of fAD, for instance, recapitulated the phenomenon of enlarged neuroprogenitor pools and fewer mature neurons^6,7^, lending weight to our mouse findings and suggesting that similar mechanisms are at play in human neural development. In summary, the developmental disturbances we observed provide a potential early fingerprint of AD mutation effect that, if validated in humans, could serve as a biomarker or a therapeutic target years before traditional prodromal AD.

### Limitations and future directions

Several limitations of the present study should be acknowledged. First, our analyses were largely conducted at the level of bulk tissue and predefined cell populations, which could obscure important cell-type-specific and cell-state-specific responses to AD mutations. The developing brain is a mosaic of diverse cell types (radial glia, intermediate progenitors, newborn neurons, oligodendrocyte precursors, microglia, etc.), each of which might be differentially affected by APP or PSEN1 mutations. A clear next step will be to apply single-cell RNA sequencing or spatial transcriptomics to embryos and juveniles carrying AD mutations. Such high-resolution approaches could identify which cell types drive the bulk changes we observed – for example, pinpointing whether NSCs themselves have altered gene expression programs or whether intermediate progenitors and neurons are primarily affected. Single-cell analysis could also reveal subtle shifts in cell proportions and uncover rare cell populations (e.g. reactive glial progenitors or dysregulated neuronal subtypes) that bulk RNA-seq cannot resolve. In our data, we saw broad signatures (upregulation of genes related to gliogenesis or neuronal migration), but the precise cellular sources of these signals remain to be mapped. A second limitation is that we focused on two AD models in mice; although they capture key aspects of mutation-driven effects, they do not encompass the full genetic landscape of AD. Other fAD mutations (in *PSEN2* or different *PSEN1* loci) and risk variants (e.g. *APOE4*, *TREM2*, *PTEN* variants affecting PI3K signaling, etc.) were not examined and could have distinct developmental impacts. Moreover, we did not include a tau-based model – tau pathology is minimal in 5xFAD and APP^NL-G-F^ mice mice, yet tau is a major mediator of neurodegeneration in AD. It remains an open question how early tau mutations or aberrant tau expression (as in some frontotemporal dementia models) might interact with developmental processes. Third, our study primarily characterized molecular and histological phenotypes; the functional consequences of these developmental changes on brain circuitry and behavior are still unknown. While young AD model mice do not exhibit overt cognitive deficits, more subtle tests of learning, memory, or sensory processing in juvenile mice could reveal whether early NSC and neuronal changes translate into functional differences. Finally, it is important to contextualize our mouse findings with human relevance. The human brain’s developmental timeline is much more protracted, and compensatory mechanisms might mitigate the impact of an *APP* or *PSEN1* mutation during childhood. Thus, any translational claims should be tempered until analogous developmental disruptions are confirmed in human studies (for example, by analyzing postmortem fetal brain tissue from individuals carrying fAD mutations, or by longitudinally observing children with familial AD risk using non-invasive imaging and biomarkers).

In conclusion, our study provides new evidence that AD-associated mutations can perturb the fundamental processes of early brain development, including neural stem cell proliferation, fate specification, and differentiation. By comparing a high-expression transgenic model (5xFAD) with a gene knock-in model (APP^NL-G-F^ mice), we disentangled the confounding influence of APP overexpression and showed that the mutations themselves – even when expressed at physiological levels – are capable of tilting the balance between neurogenesis and gliogenesis during critical developmental windows. These findings integrate with a growing body of literature suggesting that the pathogenic seeds of AD may be sown far earlier than previously appreciated (Ref: Stem cell report paper). The developmental aberrations observed here (premature stem cell differentiation, skewed neural lineage allocation, and early immune activation) could form the substrate upon which later-life pathology builds. As the field moves forward, leveraging single-cell genomics and human stem cell models will be essential to map the precise cellular and molecular cascades initiated by AD mutations in the developing brain. Unraveling these early cascades not only enriches our understanding of AD pathogenesis across the lifespan but also points to novel windows of opportunity for intervention – perhaps one day allowing us to bolster brain development in those at genetic risk for AD, long before the inexorable decline into neurodegeneration begins.

## Materials and Methods

### Animals

All mouse experiments were conducted in accordance with the guidelines of the Swedish Board of Agriculture (ethical permit 12570-2021) and were approved by the Karolinska Institutet Animal Care Committee. The mouse lines (C57Bl6/J or APP^NL-G-F^) were bred with C57Bl6/J so that the same genetic background allowed us to safely compare the different conditions.

### Tissue processing

At the end of the survival period, the pregnant dam were sacrificed using carbon dioxide chambers, and postnatal pups were quickly decapitalized. Brains were dissected in cold PBS, then were used directly for cell culture, or further postfixed in 4% PFA in PBS at 4°C overnight and cryoprotected in 30% sucrose for at least 48h. For PFA fixed tissues, brains were embedded in Tissue-Tek OCT compound (Sakura), and were sectioned coronally to 16– 20µm thickness. Sections were collected accordingly to stereological principles and stored at −80°C until further use.

### RNAseq

RNAseq experiments were performed as we previously described^32^. Briefly, total RNA was extracted from the cortex or hippocampi of each genotype at indicated timepoints, by using QIAGEN RNeasy Mini kit, following the supplier’s instructions. Extracted RNA was sent for BEA core facility at Karolinska Institute for the measurement of integrity (RIN) and concentration, by using Agilent RNA 600 Nano kit and the Agilent 2000 Bioanalyzer system (Agilent Technologies, Sweden). RNA-sequencing was performed by National Genomics Infrastructure (NGI) at Science for Life Laboratory (Stockholm, Sweden), using the S4 flow cell and sequenced on a NovaSeq 6000 (Illumina Inc, San Diego, CA) with read length of 2 x 150 bp.

### Cell culture

Dissected embryonic forebrains were dissociated into single-cell suspension and neurosphere cultures were established as we described before^33,34^. After counting the cell number from each tissue, cells isolated from each animal were plated in T25 flasks. Primary neurospheres were harvested after 7 days in culture for quantification, then were dissociated into single cells for passage or differentiation. Approximately 100,000 cells per animal were used for new generation of neurospheres or for differentiation assay, and quantifications for next generation of neurospheres will be quantified 7 days after culture. For differentiation assay, primary neurospheres were dissociated, single cells were seeded in poly-L-lysine-coated chamber slides (Sigma) at approximately 10,000 cells/well for E13.5, and 20,000 cells/well for E18.5, with removal of growth factors. 14 days after culture, differentiated cells were fixed by 4% PFA, then subjected for immunofluorescence assay.

### Immunohistochemistry and Immunocytochemistry

Tissue sections or PFA fixed differentiated cells were prcessed as we previously described for normal donkey serum and permebalization^33,34^. Primary antibodies were incubated at room temperature (RT) overnight and secondary antibodies were incubated for 1h at RT. Secondary antibodies were used accordingly. Counterstaining was performed with DAPI (1:5000) in PBS and sections were coverslipped with Vectashield mounting media. Full details of the primary antibodies used are reported in Table 2.

### Imaging and quantification

Confocal representative images were acquired using Leica confocol microscope set up. For the quantification, the regions of interest were analyzed either manaually or via Qupath. The tissue quantification was performed on at least 3 sections per segment and area. For each experimental group and marker, 3–9 animals were analyzed.

### Statistics

For image quantification, Student’s T-test for comparing two groups and One-way Annova for more than two groups’ comparisons were performed by statistical analysis software Prism version 10. *P<0.05, **P<0.01, ***P<0.001. For each experimental group and marker staining, 3–9 animals were analyzed.

### RNAseq analysis

The data preparation was done in BEA core facility as previously described (Wang et al., 2025). High-quality RNA-Seq paired reads were mapped to the mouse reference genome using STAR aligner ^35^. Differential gene expression analysis was performed on featured gene counts with R/Bioconductor EdgeR software ^36^. Adjusted p-value of < 0.05 was used to determine differentially expressed genes (DEGs). Three to four biological replicates were included for all comparative analyses. Biofunction and pathway analysis was performed using “clusterProfiler” package ^37^ based on GO, KEGG ^38^ and the MSigDB ^39^. Gene set enrichment was evaluated by normalized enrichment score (NES) as calculated by actual ES divided by the mean (ES against all permutations of the data set). Detailed information regarding KEGG pathways was visualized using Pathview ^40^.

Single-cell RNA sequencing (scRNAseq) data was obtained from Di Bella et al.^21^ and processed using the Seurat R package (version 4.4.0) as we previously decribed^41^. NormalizeData function was used for data normalization. For dimensionality reduction, the original embeddings from the study^21^ were applied. Differential gene expression analysis for different cell types and timepoints was performed using default parameters of FindAllMarkers function in Seurat.

## Data availability

RNAseq datasets produced in this manuscript will be available in the Gene Expression Omnibus after the manuscript has been published in peer-review journal. The publicly available data used in this study is available at: Di Bella: https://www.ncbi.nlm.nih.gov/geo/query/acc.cgi?acc=GSE153164

## Code availability

Software used for analysis is public and described in detail in the Methods section. Raw scripts and code are available upon publication of this manuscript.

## Acknowledgements

We acknowledge Dr. Per Nilsson and Dr. Cecilia Dominguez for organizing the animals in the division, Dr. Luana Naina for supporting embryonic tissue dissection. Mr. Ling Zhang for supporting neurosphere assay.

